# Applied phenomics and genomics for improving barley yellow dwarf resistance in winter wheat

**DOI:** 10.1101/2022.01.05.475073

**Authors:** Paula Silva, Byron Evers, Alexandria Kieffaber, Xu Wang, Richard Brown, Liangliang Gao, Allan Fritz, Jared Crain, Jesse Poland

**Affiliations:** Department of Plant Pathology, College of Agriculture, Kansas State University, Manhattan, Kansas, 66506; Programa Nacional de Cultivos de Secano, Instituto Nacional de Investigación Agropecuaria (INIA), Estación Experimental La Estanzuela, Colonia, Uruguay, 70006; Department of Agricultural and Biological Engineering, University of Florida, IFAS Gulf Coast Research and Education Center, Wimauma, Florida, 33598; Department of Agronomy, College of Agriculture, Kansas State University, Manhattan, Kansas, 66506; King Abdullah University of Science and Technology, Thuwal, Saudi Arabia

**Keywords:** Barley yellow dwarf (BYD), High-throughput Phenotyping (HTP), *Triticum aestivum*, Virus, Resistance, Tolerance, Genome-wide Association Mapping (GWAS), Genomic Selection (GS)

## Abstract

Barley yellow dwarf (BYD) is one of the major viral diseases of cereals. Phenotyping BYD in wheat is extremely challenging due to similarities to other biotic and abiotic stresses. Breeding for resistance is additionally challenging as the wheat primary germplasm pool lacks genetic resistance, with most of the few resistance genes named to date originating from a wild relative species. The objectives of this study were to, i) evaluate the use of high-throughput phenotyping (HTP) from unmanned aerial systems to improve BYD assessment and selection, ii) identify genomic regions associated with BYD resistance, and iii) evaluate genomic prediction models ability to predict BYD resistance. Up to 107 wheat lines were phenotyped during each of five field seasons under both insecticide treated and untreated plots. Across all seasons, BYD severity was lower with the insecticide treatment and plant height (PTHTM) and grain yield (GY) showed increased values relative to untreated entries. Only 9.2% of the lines were positive for the presence of the translocated segment carrying resistance gene *Bdv2* on chromosome 7DL. Despite the low frequency, this region was identified through association mapping. Furthermore, we mapped a potentially novel genomic region for resistance on chromosome 5AS. Given the variable heritability of the trait (0.211 – 0.806), we obtained relatively good predictive ability for BYD severity ranging between 0.06 – 0.26. Including *Bdv2* on the predictive model had a large effect for predicting BYD but almost no effect for PTHTM and GY. This study was the first attempt to characterize BYD using field-HTP and apply GS to predict the disease severity. These methods have the potential to improve BYD characterization and identifying new sources of resistance will be crucial for delivering BYD resistant germplasm.

## Introduction

Wheat (*Triticum aestivum* L.) is one of the most essential food crops in the world and is constantly threatened by several biotic stresses (Savary *et al*. 2019). Among the most important viral stresses is barley yellow dwarf (BYD). This disease is widespread across the world, caused by viruses and transmitted by aphids (Shah *et al*. 2012), and can cause significant yield reductions in susceptible cultivars. In Kansas, BYD is the fourth most significant wheat disease in terms of average estimated yield losses with an average yield loss of approximately 1% estimated over the past 20 years (Hollandbeck *et al*. 2019), equivalent to a loss of more than $10 million per year. However, yield losses are highly variable ranging from 5% to 80% in a single field depending on the environment, management practices, the host, and the genetic background, (Miller and Rasochová 1997; Perry *et al*. 2000; Gaunce and Bockus 2015). Moreover, the wide host range and the complex lifestyle of its vectors make BYD extremely difficult to manage, and different management strategies (e.g., planting date and control of vector populations) are inconsistent depending on climate and location (Bockus *et al*. 2016). Thus, in many production environments, particularly in the Central and Eastern regions of Kansas, BYD is often the most economically impactful disease.

Barley yellow dwarf disease symptoms are highly variable depending on the crop, variety, time, and developmental stage when the infection occurs, aphid pressure, and environmental conditions (Shah *et al*. 2012; Choudhury *et al*. 2019b). BYD characterization in the field is extremely challenging as the symptoms can easily be confused with other viral disease symptoms such as wheat streak mosaic virus symptoms, nutrient deficiencies, or environmental stresses like waterlogging (Shah *et al*. 2012). Typical BYD symptoms can be observed at all levels of plant organization – leaf, roots, and flowers. Leaf discoloration in shades of yellow, red, or purple, specifically starting at the tip of the leaf and spreading from the margins toward the base is common as well as a reduction in chlorophyll content (Jensen and Van Sambeek 1972; D’arcy 1995). Often the entire plant visually appears stunted or dwarfed from a reduction in biomass by reducing tiller numbers. Spike grain yield is decreased through a reduction in kernels per spike and kernel weight which also affects grain quality (Riedell *et al*. 2003; Choudhury *et al*. 2019b). Quality can be further reduced by a reduction in starch content (Peiris *et al*. 2019). Below ground effects of BYD have also been reported including reduced root growth (Riedell *et al*. 2003).

Currently, there is no simple solution to control BYD (Walls *et al*. 2019), however, the use of genetic resistance and tolerance is the most appealing and cost-effective option to control this disease (Comeau and Haber 2002; Choudhury *et al*. 2017; 2019b). Resistance and tolerance could be different genetic mechanisms, namely stopping virus replication and minimizing disease symptoms respectively, but within this paper all mention of resistance includes both genetic resistance and tolerance. Breeding strategies involving genetic resistance can target either the aphids or the virus. Resistance to aphids can be achieved by three different strategies, antixenosis, antibiosis, or tolerance (Girvin *et al*. 2017). To date, most breeding efforts have been directed to the identification of viral tolerance, also known as ‘field resistance’, that refers to the ability of the plant to yield under BYD infection and is associated with a reduction of symptoms of infection independent of the virus titer (Foresman *et al*. 2016). Field resistance has been reported to be polygenic, falling under the quantitative resistance class, where several genes with very small effects control the resistance response (Qualset *et al*. 1973, Cisar *et al*. 1982; Ayala *et al*. 2002; Choudhury *et al*. 2019a; c).

Presently, no major gene conferring immunity or a strong resistant phenotype to BYD has been identified in bread wheat, and only four resistance genes have been described for BYD. Located on chromosome 7DS, *Bdv1* is the only gene described from the primary pool of wheat and was originally identified in the wheat cultivar ‘Anza’ (Qualset *et al*. 1984; Singh *et al*. 1993). This gene provides resistance to some but not all the viruses that cause BYD (Ayala-Navarrete and Larkin 2011). The other three named genes were all introduced into wheat through wide crossing from intermediate wheatgrass (*Thinopyrum intermedium*) (Ayala *et al*. 2001; Zhang *et al*. 2009). *Bdv2* and *Bdv3* are both located on a translocation segment on wheat chromosome 7DL (Brettell *et al*. 1988; Sharma *et al*. 1995), while *Bdv4* is located on a translocation segment on chromosome 2D (Larkin *et al*. 1995; Lin *et al*. 2007). *Bdv2* was the first gene successfully introgressed in wheat breeding programs from the tertiary gene pool for BYD resistance (Banks *et al*. 1995) and deployed into varieties.

In addition to the four known resistance genes, other genomic regions associated with BYD resistance have been identified through genetic mapping. These regions have been described on nearly all wheat chromosomes but have not been genetically characterized (Ayala *et al*. 2002; Jarošová *et al*. 2016; Choudhury *et al*. 2019a; b; c). Moreover, two recent studies have reported that some of these new genomic regions display additive effects (Choudhury *et al*. 2019a; b). Additive genetic effects had already been reported in lines combining *Bdv2* and *Bdv4* (Jahier *et al*. 2009).

Taken together, research indicates that resistance genes to BYD in wheat are rare. With a lack of major genes and difficulty to characterize resistance in the wheat pool likely due to the polygenic nature of many small effect loci, identifying resistance has been limited. Nevertheless, breeding programs have devoted large efforts for breeding BYD resistance due to the economic importance of this disease, with some of the greatest success coming from wide crosses to the tertiary gene pool.

Breeding for BYD resistance can be improved by applying strategies for more effective evaluation and utilization of the identified resistance. To get a better understanding of BYD and its quantitative nature, consistent and high-throughput methods are needed for the identification of resistant wheat lines for large-scale selection in breeding programs (Aradottir and Crespo-Herrera 2021). Effective selection on the quantitative resistance with low heritability can be aided by the high-throughput genotyping, high-throughput phenotyping (HTP), or a combination of both.

Access to high-density genetic markers at a very low-cost, owing to the rapid developments in DNA sequencing, have enabled breeding programs to apply molecular breeding for quantitative traits. Genomic selection (GS) is a powerful tool to breed for quantitative traits with complex genetic architecture and low heritability (e.g., yield, quality, and diseases such as Fusarium head blight), because it has greater power to capture loci with small effect compared with other marker-assisted selection strategies (Meuwissen *et al*. 2001; Poland and Rutkoski 2016). In addition to molecular data, HTP using unmanned aerial systems (UAS), or ground-based sensors is providing high density phenotypic data that can be incorporated into breeding programs to increase genetic gain (Haghighattalab *et al*. 2016; Crain *et al*. 2018; Wang *et al*. 2020). Using precision phenotyping for disease scoring can improve the capacity for rapid and non-biased evaluation of large field-scale numbers of entries (Poland and Nelson 2011). Taken together improvements in genomics and phenomics have the potential to aid breeding progress for BYD resistance.

In an effort to accelerate the development of resistant lines, we combined high throughput genotyping and phenotyping to assess BYD severity in a large panel of elite wheat lines. We evaluated the potential of HTP data to accurately assess BYD severity as well as identify genetic regions associated with BYD resistance and inform whole genome prediction to identify resistant lines.

## Materials and Methods

### Plant Material

A total of 381 different wheat genotypes were characterized for BYD resistance, including 30 wheat cultivars and 351 advanced breeding lines in field nurseries over five years (Table S1). In each nursery, an unbalanced set of 52 – 107 wheat entries were evaluated including both cultivars and breeding lines (Table 1). The BYD susceptible cultivar ‘Art’ and BYD resistant cultivar ‘Everest’ were included in all the nurseries (seasons) as checks.

**Table 1:** Field experimental details for the five wheat nurseries

| Season | 2015 – 2016 | 2016 – 2017 | 2017 – 2018 | 2018 – 2019 | 2019 – 2020 |
| --- | --- | --- | --- | --- | --- |
| Location | Rocky Ford farm |  | Ashland Bottoms farm |  |  |
|  | 39°13'45.60" N, 96°34'41.21" W |  | 39°07'53.76" N, 96°37'05.20" W |  |  |
| Planting Date | Sep. 17, 2015 | Sep. 12, 2016 | Sep. 19, 2017 | Sep. 17, 2018 | Sep. 17, 2019 |
| Number of Entries | 68 | 52 | 81 | 81 | 107 |
| Number of Plots | 504 | 360 | 400 | 392 | 684 |
| Field Design | split-plot with insecticide treatment as major factor effect and wheat genotype as secondary factor |  |  |  |  |
| Replications | 3 | 3 | 2 | 2 | 2 |
| Plot Size | 6 rows plots - 1.5 m × 2.4 m |  |  |  |  |
| BYD Evaluation | April 28, 2016 | May 12, 2017 | May 19, 2018 | May 13, 2019 | May 19, 2020 |
| Harvesting Date | June 20, 2016 | June 19, 2017 | June 23, 2018 | June 28, 2019 | June 25, 2020 |

### Field Experiments

Nurseries for BYD field-screening were conducted during five consecutive wheat seasons (2015 – 2016 to 2019 – 2020) (Table 1). Seasons 2015 – 16 and 2016 – 17 were conducted at Kansas State University (KSU) Rocky Ford experimental station (39°13′45.60″ N, 96°34′41.21″ W), while the 2017 – 18, 2018 – 19, and 2019 – 20 nurseries were planted at KSU Ashland Bottoms experimental station (39°07′53.76″ N, 96°37′05.20″ W). The nurseries were established for natural infections by planting about three weeks earlier than the normal planting window in mid-September. The susceptible cultivar ‘Art’ was planted as a spreader plot in the borders and as a control check plot also with the resistant cultivar ‘Everest’. The experimental unit was 1.5m × 2.4m with a six-row plot on 20cm row spacing.

A split-plot field design with two or three replications was used where the main plot was insecticide treatment, and the split plot was the wheat genotype. Three replications were used for proof of concept during the first two seasons but then two replications were chosen as a balance of space and number of entries for the following seasons. For the treated replications the seed were treated at planting with Gaucho XT (combination of insecticide and fungicide) at a rate of 0.22 ml/100g of seed, followed with foliar insecticide applications starting from approximately 2 – 3 weeks after planting through heading. Depending on field conditions, spray treatments were conducted every 14 – 21 days if average air temperatures remained above 10°C. Foliar insecticides were applied to the treated replications in a spray volume of 280.5 L/ha using a Bowman MudMaster plot sprayer equipped with TeeJet Turbo TwinJet tips. Insecticide applications consisted of a rotation of Warrior II, Lorsban, and Mustang Max at rates of 0.14L/ha, 1.17L/ha, and 0.29L/ha, respectively. For the control insecticide treatment (untreated), the seed were treated with Raxil MD (fungicide) at a rate of 0.28ml/100g of seed, and no foliar insecticide applications were applied. Foliar fungicide Nexicor was applied to the whole experiment at a rate of 0.73L/ha, at both planting and heading, to control all other diseases so the main disease pressure was focused on BYD.

### Phenotypic Data

Individual plots were assessed for i) BYD severity characterized as the typical visual symptoms of yellowing or purpling on leaves using a 0 – 100% visual scale, determined directly after spike emergence by recording the proportion of the plot exhibiting the symptoms (Table 1), ii) manual plant height (PTHT_M_, meters), and iii) grain yield (GY, tons/ha). Experimental plots were harvested using a Kincaid 8XP plot combine (Kincaid Manufacturing., Haven, KS, USA). Grain weight, grain moisture and test weight measurements for each plot was recorded using a Harvest Master Classic GrainGage and Mirus harvest software (Juniper Systems, Logan, UT, USA). Visual phenotypic assessment was recorded using the Field Book phenoapp (Rife and Poland 2014).

### High-Throughput Phenotyping

To compliment the manually recorded phenotypic data, we applied HTP using a ground-based proximal sensing platform or an UAS (Table 2). Seasons 2015 – 16 and 2016 – 17 were characterized by the ground platform as described in Barker *et al*. (2016) and Wang *et al*. (2018). For the other three seasons, we used a quadcopter DJI Matrice 100 (DJI, Shenzhen, China) carrying a MicaSense RedEdge-M multispectral camera (MicaSense Inc., United States). The HTP data was collected on multiple dates throughout the growth cycle from stem elongation to ripening (GS 30 – 90; Zadoks *et al*. 1974) (Table 2). Flight plans were created using CSIRO mission planner application and missions were executed using the Litchi Mobile App (VC Technology Ltd., UK, https://uavmissionplanner.netlify.app/) for DJI Matrice100. The aerial image overlap rate between two geospatially adjacent images was set to 80% both sequentially and laterally to ensure optimal orthomosaic photo stitching quality. All UAS flights were set at 20m above ground level at 2m/s and conducted within two hours of solar noon. To improve the geospatial accuracy of orthomosaic images, white square tiles with a dimension of 0.30m × 0.30m were used as ground control points and uniformly distributed in the field experiment before image acquisition and surveyed to cm-level resolution using the Emlid REACH RS+ Real-Time Kinematic Global Navigation Satellite System unit (Emlid Ltd., HongKong, China).

**Table 2.** Dates of high-throughput phenotypic data collection and details of image acquisition in the five wheat nurseries screened for BYD, Kansas, USA (2015-2020).

| Season | 2015 – 2016 | 2016 – 2017 | 2017 – 2018 | 2018 – 2019 | 2019 – 2020 |
| --- | --- | --- | --- | --- | --- |
| UAS Platform | PheMU |  | DJI Matrice 100 |  |  |
| Imaging Sensor | multiple digital single-lens reflex (DSLR) cameras |  | MicaSense RedEdge-M |  |  |
| Flight/Pass speed | 0.3–0.5 m/s |  | 2 m/s |  |  |
| Flight Dates | 2016-03-31<br>2016-04-07<br>2016-04-14<br>2016-05-06 |  |  | 2019-04-01 |  |
|  |  |  |  | 2019-04-09 |  |
|  |  | 2017-03-28 |  | 2019-04-19 |  |
|  |  | 2017-04-13 | 2018-03-30 | 2019-04-26 | 2020-03-20 |
|  |  | 2017-05-01 | 2018-04-04 | 2019-05-02 | 2020-04-11 |
|  |  | 2017-05-09 | 2018-04-12 | 2019-05-10 | 2020-04-23 |
|  |  | 2017-05-21 | 2018-04-19 | 2019-05-15 | 2020-05-03 |
|  |  | 2017-05-23 | 2018-04-23 | 2019-05-23 | 2020-05-19 |
|  |  | 2017-05-30 | 2018-05-16 | 2019-05-31 | 2020-06-05 |
|  |  | 2017-06-05 | 2018-06-13 | 2019-06-05 | 2020-06-11 |
|  |  | 2017-06-13 |  | 2019-06-12 |  |
|  |  |  |  | 2019-06-17 |  |
| Flight/Pass altitude | 0.5 m above the canopy |  | 20 m AGL |  |  |
| In-Air Flight Duration | NA |  | ~ 11–14 min |  |  |

An automated image processing pipeline (Wang *et al*. 2020) was used to generate the orthomosaics and extract plot-level plant height (PTHT_D_ (m), Singh *et al*. 2019) and the normalized difference vegetation index (NDVI) (Rouse *et al*. 1974), calculated as:

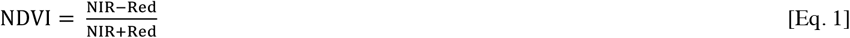

where NIR and Red are the near-infrared and red bands of the multispectral images and NDVI is the output image. Both traits were selected based on potential BYD characterization where the most typical BYD symptoms include chlorosis and stunting of the plants, thus, influencing NDVI and PTHT.

### Statistical Data Analyses

First, the adjusted mean best linear unbiased estimator (BLUE) was calculated for each entry for all the different traits for each season (Table S1), using the following model:

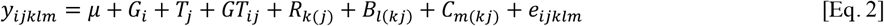

where *y_ijklm_* is the phenotype for the trait of interest, *μ* is the overall mean, *G_i_* is the fixed effect of the *i^th^* entry (genotype), *T_j_* is the fixed effect of the *j^th^* insecticide treatment, *GT_ij_* is the fixed effect of the interaction between the *i^th^* entry and the *j^th^* insecticide treatment (genotype by treatment effect), *R*_*k*(*j*)_ is the random effect of the *k^th^* replication nested within the *j^th^* insecticide treatment and distributed as iid 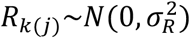, *B*_*l*(*kj*)_ is the random effect of the *l^th^* row nested within the *k^th^* replication and *j^th^* treatment distributed as iid 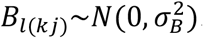, *C*_*m*(*kj*)_ is the random effect of the *m^th^* column nested within the *k^th^* replication and *j^th^* treatment and assumed distributed as iid 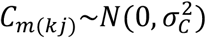, and *e_ijklm_* is the residual for the *ijklm^th^* plot and distributed as iid 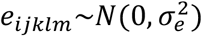. The ‘lme4’ R package (Bates *et al*. 2014) was used for fitting the models.

The BLUEs were used to inspect trait distributions and to calculate Pearson correlations between all traits. In addition, BLUE values were used to calculate the reduction in GY for each entry as the difference of GY between the untreated and insecticide treated main plots. This variable reflects the level of BYD resistance of each entry, and it was used to perform GWAS and GS analyses.

For NDVI and PTHT_D_, the plot-level observed values extracted for the different phenotypic dates were fitted to a logistic non-linear regression model (Fox and Weisberg 2011) as,

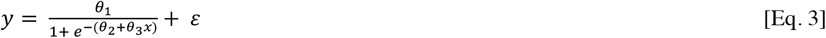

where is *y* the phenotype for the trait of interest at the time-point *x* measured as days after January 1, *θ*_1_ is the maximum value (upper asymptote) represented by the final PTHT or maximum achieved NDVI, *θ*_2_ is the inflection point that represents the greatest rate of change in the growth curve, either senescence for NDVI or height of growth, *θ*_3_ is the lag phase or onset of senescence or growth rate from time *x* where x is the calendar day of the year since January 1, and *ε* is the residual error (Figure S1). The “nlme” R package was used for model fitting (Pinheiro *et al*. 2015). The model parameters obtained for each trait (*θ*_1*NDVI*_, *θ*_2*NDVI*_, *θ*_3*NDVI*_, *θ*_1*PTHT_D_*_, *θ*_2*PTHT_D_*_, and *θ*_3*PTHT_D_*_) were used in addition to the other phenotypic traits to calculate BLUEs, distributions, correlations, and BLUPs.

Secondly, we used a mixed linear model to calculate the best linear unbiased predictors (BLUPs) for each entry in each nursery (season) (Table S1), using the same model as described in equation 2 but defining *G_i_*, *T_j_*, and *GT_ij_* as random effects. BLUPs were used because of the unbalanced nature of the data (not all lines were evaluated in all the seasons). The BLUPs calculated for each season were then combined for GWAS and GS. Furthermore, we calculated broad-sense heritability on a line-mean basis by splitting the data based on whole plot treatment for insecticide treatments as:

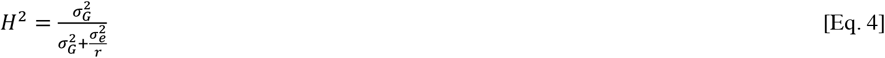

where 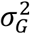 is the genotypic variance, 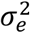 is the residual error variance, and *r* is the number of replications.

### Genotypic Data

A total of 346 wheat entries were genotyped using genotyping-by-sequencing (GBS) (Poland *et al*. 2012) and sequenced on an Illumina Hi Seq2000. Single nucleotide polymorphisms (SNPs) were called using Tassel GBSv2 pipeline (Glaubitz *et al*. 2014) and anchored to the Chinese Spring genome assembly v1.0 (Appels *et al*. 2018). SNP markers with minor allele frequency < 0.01, missing data > 85%, or heterozygosity > 15% were removed from the analysis. After filtering, we retained 29,480 SNPs markers that were used to investigate the population structure through principal component analysis (PCA), genome-wide association analysis (GWAS), and GS. In addition, GBS data was used to run a bioinformatics pipeline to predict the presence or absence of the translocated segment on chromomere 7DL carrying the *Bdv2* gene for each entry (Table S1). The prediction was done based on a modified alien predict pipeline (Gao *et al*. 2021). Briefly, alien or wheat specific tags were counted in the 7DL region and tabulated using a training set of cultivars or lines that are known to be *Bdv2* positive and negative. A simple classification was done based on alien to wheat tag counts ratios.

### Genome-Wide Association Analysis

The GWAS analysis was performed with a mixed linear model implemented in the ‘GAPIT’ R package (Lipka *et al*. 2012) that includes principal components to account for population structure as fixed effects and the individuals to explains familial relatedness as random effects,

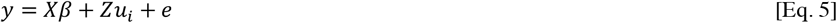

where *y* is the vector of phenotypic BLUPs, *X* and *Z* are the incidence matrix of *β* and *u_i_*, respectively, with *u_i_* assumed 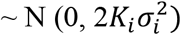 where *K* is the individual kinship matrix, and *e* is the vector of random residual effects with 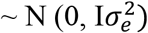, where *I* is the identity matrix and 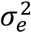 is the unknown residual variance. The false discovery rate correction with an experimental significance level value of 0.01 was used to assess marker-trait associations. Manhattan plots were generated with ‘CMplot’ package in R software (Yin 2020). PCA using GBS-SNPs was performed in R language. Eigenvalues and eigenvectors were computed with ‘e’ function using ‘A.mat’ function and the ‘mean’ imputation method of ‘rrBLUP’ package (Endelman, 2011). To declare a quantitative trait locus (QTL) we considered only the regions having several SNP markers in linkage disequilibrium, clearly showing a peak. We did not consider regions with a single SNP above the significant threshold as a QTL.

### Genomic Selection

Using data from the five seasons, GS models using the genomic best linear unbiased predictor (G-BLUP) were developed to assess predictive ability. A five-fold cross-validation method was used to assess model accuracy where the data set was split into five sets based on season, with four seasons forming the training set and the fifth season serving as prediction set. This process was repeated until all seasons were predicted. Along with predicting all other seasons from each season, a model was evaluated with a leave-two-out cross-validation strategy. This strategy was used to get a better mix of years with and without disease incidence, where the training population consisted of three seasons, and the remaining two seasons were predicted from the combined training population. The GS model was fitted with the training population using ‘rrBLUP’ *kin.blup* function (Endelman 2011), the GS model equation was,

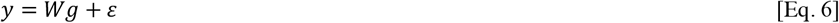

where *y* is a vector of phenotypic BLUPs, *W* is the design matrix of *g, g* is the vector of genotypic values 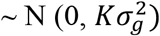 and *ε* is the vector of residual errors (Endelman 2011). Predictive ability was assessed using Pearson’s correlation (*r*) between the predicted value (G-BLUP) and the BLUP for the respective phenotype. In addition, for both GS strategies we also tested the effect of adding the genotype of the *Bdv2* loci as a fixed effect cofactor, using the model,

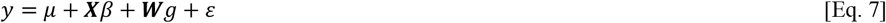

which combines parameters described in equation 6 and *X* is the matrix (*n x* 1) of individual observation for presence or absence of *Bdv2* and *β* is the fixed effect for the *Bdv2* measurements.

## Results

### Phenotypic Data

We analyzed five years of BYD field-screening nurseries (seasons 2015-16 to 2019-20) characterizing a total of 381 wheat lines. The disease pressure and the expression of BYD associated symptoms varied each season, however, we were able to observe a significant effect of the insecticide treatment in all seasons (Figure 1). Across all seasons, BYD symptoms were lower on the insecticide treated plots and both PTHT_M_ and GY increased compared to the non-treated control. Season 2016-17 had the most conducive conditions for BYD screening, resulting in high average severity and a larger difference between mean values for the treated vs untreated plots for all the collected traits (Figure 1). There was general consistency in order across all seasons with the susceptible check ‘Art’ ranked among the highest in BYD severity (Figure S2).

**Figure 1.**
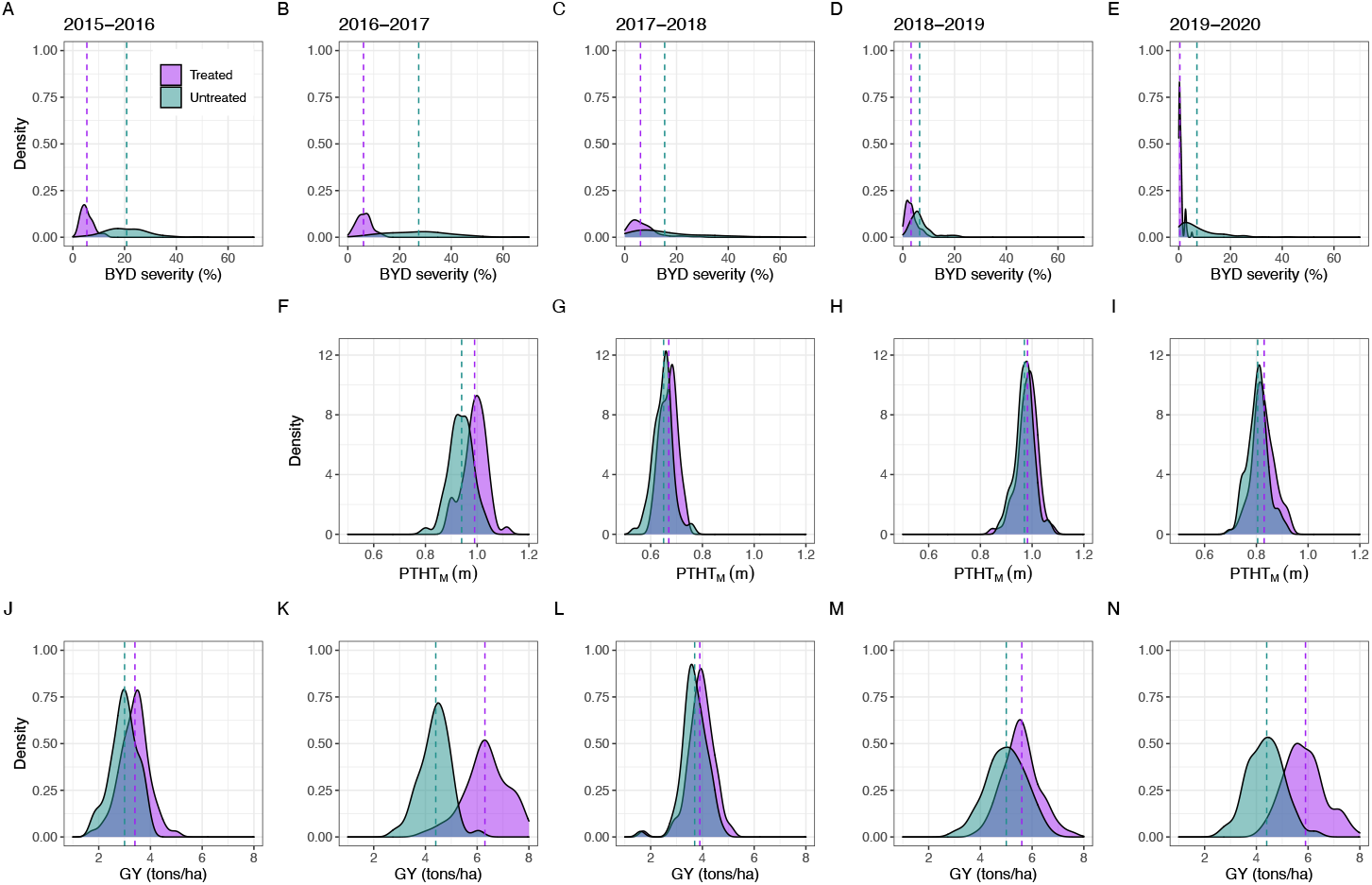
Adjusted phenotypic values for the traits collected manually for five different field seasons (2015-2016 to 2019-2020). A-E): barley yellow dwarf (BYD) severity (%) characterized as the typical visual symptoms of yellowing/purpling on leaves using a 0 – 100% visual scale, FI) manual plant height/stunting (PTHT_M_) (meters), note that the trait was not recorded for the 2015 – 2016 season, and J-N) grain yield (GY) (tons/ha). Insecticide-treated and untreated replications are represented by purple and green, respectively. The dashed line represents the mean value for the trait in each treatment

Phenotypic correlations between the traits showed a negative correlation between BYD and GY for all the seasons and a negative or no correlation between BYD and PTHT_M_ (Figure S3). The same correlation trends were observed under insecticide treated and untreated plots. Broad-sense heritability was moderate to high for all the traits, ranging between 0.21 and 0.79 for the insecticide treated plots and between 0.41 and 0.84 for the untreated plots. Across all traits, the untreated insecticide replications showed higher *H*^2^ values, with season 2016 – 17 showing the highest values (Figure 2).

**Figure 2.**
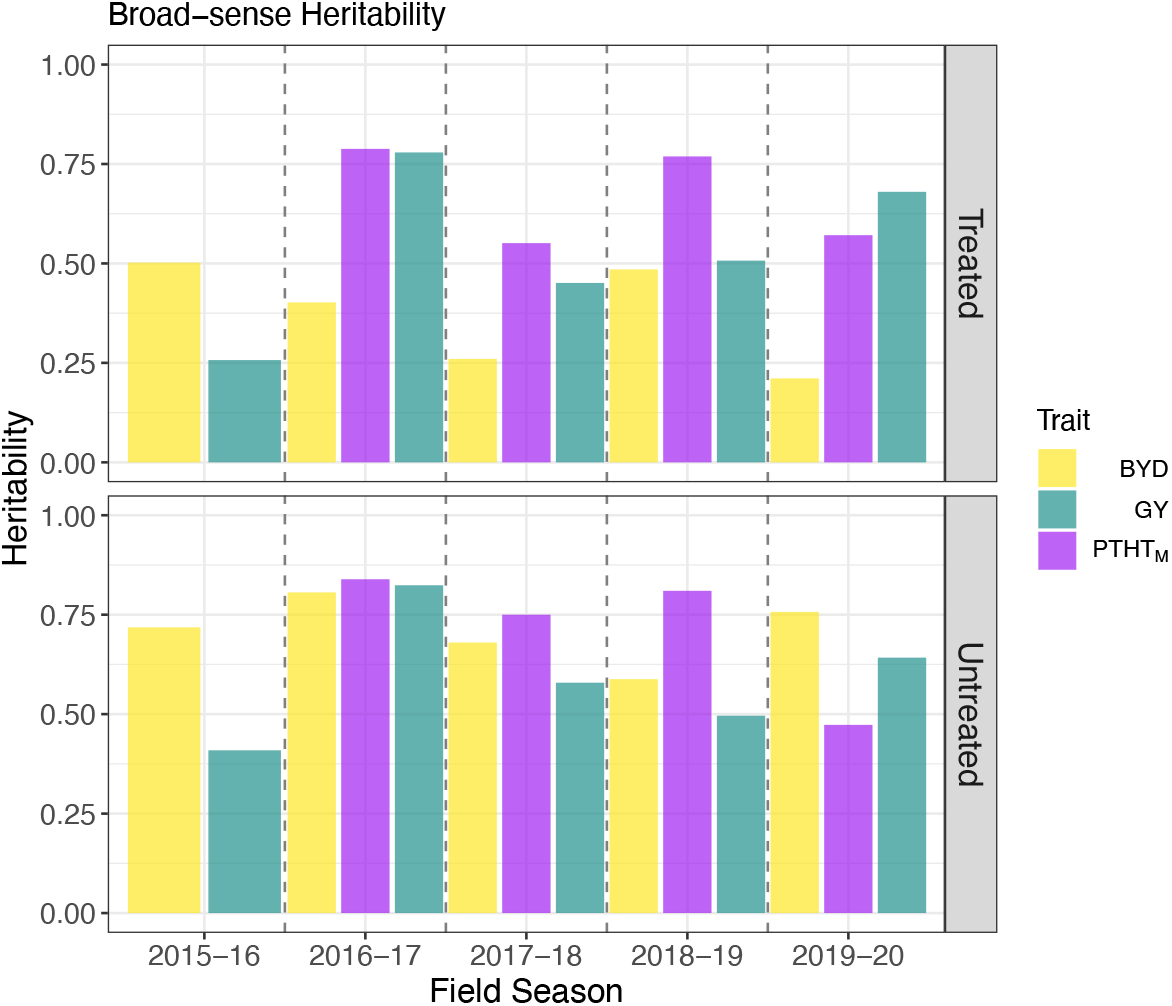
Broad-sense heritability of wheat phenotypic traits collected manually, including visual barley yellow dwarf (BYD) score, plant height (PTHT_M_) and grain yield (GY) during five different field seasons under two insecticide treatments.

For the HTP data collected (Table 2), we obtained three different parameters (*θ*_1_, *θ*_2_, and *θ*_3_) for both PTHT_D_ and NDVI after fitting a logistic regression model using the data collected during the experiments (2015-16 season data was not included due to lack of data quality) (Figure S1). Correlations between these parameters and the phenotypic traits collected manually were different for all the traits (Figure S3). For the insecticide untreated plots, BYD resulted in a negative correlation with *θ*_2*NDVI*_ and a positive correlation with *θ*_3*NDVI*_, in most of the field seasons. We did not find a clear correlation pattern between BYD and *PTHT_D_*. For PTHT_M_ we detected a positive correlation with *θ*_1*PTHT_D_*_ across all seasons, and for GY we observed a positive correlation with *θ*_1*NDVI*_ and *θ*_2*NDVI*_, and a negative correlation with *θ*_3*NDVI*_ (Figure S3).

### Prediction of *Bdv2* Resistance Gene

We used GBS data to genotype the *Bdv2* resistance gene located on a translocation segment from intermediate wheatgrass on chromosome 7DL of bread wheat. In total, 33 of the 346 wheat lines carried the *Th. intermedium* chromosomal translocation with *Bdv2* (Table S1). Interestingly, 28 of these *Bdv2* lines belonged to the same breeding cycle, entering the advanced yield nursery stage of the KSU breeding program in the 2017 – 18 season. Furthermore, only 7 pedigrees are represented within the 28 *Bdv2* entries, meaning that these lines are highly related. The remaining 5 *Bdv2* lines were distributed in 2015 – 16 (n=3), 2018 – 19 (n=1), and 2019 – 20 (n=1), and none of the lines from the season 2016 – 17 had the presence of *Bdv2* (Table S1).

### Population Structure

We studied the population structure of 346 wheat lines using 29,480 GBS-derived SNP markers. The PCA did not reveal a strong pattern of population structure (Figure 3). Moreover, the variation explained by the first two principal components (4 and 3%, respectively) also supports the hypothesis of minimal population structure within a single breeding program. We observed that most of the wheat cultivars released by KSU breeding program were located outside the cluster grouping all the breeding lines (Figure 3A). Lines with the presence of *Bdv2* clustered together (Figure 3B), likely due to a related pedigree to the original source, and we did not identify any evident pattern for BYD severity associated with the population structure (Figure 3C).

**Figure 3.**
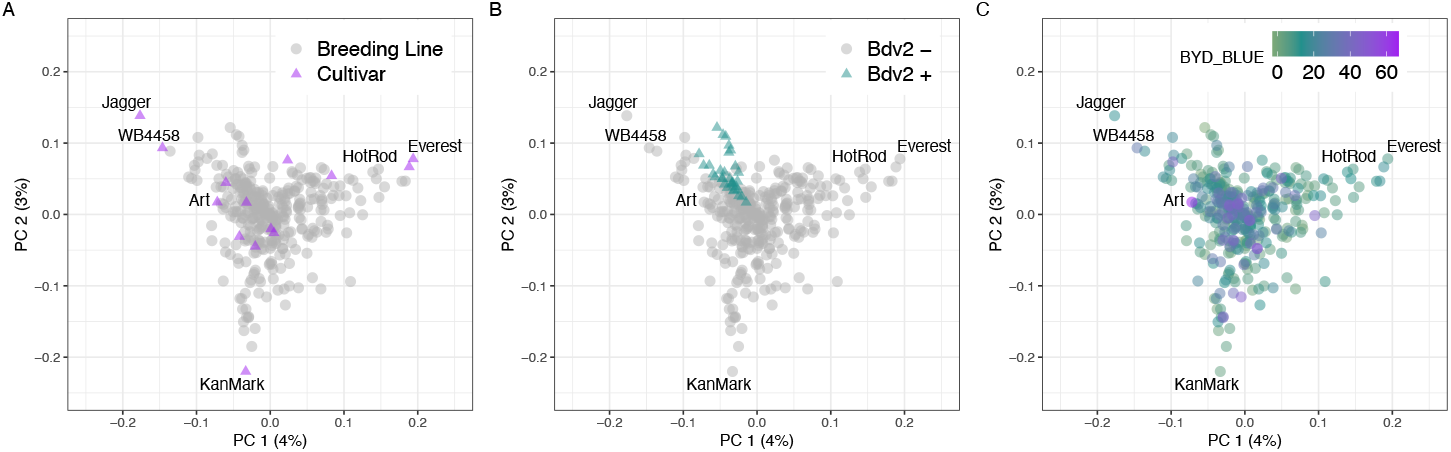
Scatterplot of the first two principal component axis, made from principal component analysis on the marker matrix, n = 357 wheat lines, markers = 29,480. Each data point represents an individual wheat line that is color-coded by A) breeding status, B) prediction of *Bdv2* presence/absence, and C) adjusted mean for BYD severity (BYD BLUE) scored visually. Total variance explained by each principal component (PC) is listed on the axis.

### Genome-Wide Association Analysis

To investigate the genetic architecture of BYD we performed GWAS analyses for all collected traits using the BLUP values for 346 lines and 29,480 SNP markers. The first two principal components from PCA and the kinship matrix were included in the mixed model to account for population structure and genetic relatedness. We found significant marker-trait associations for BYD severity on chromosomes 5AS, 7AL, and 7DL (Figure 4A). The highest peak was observed on the proximal end of chromosome 7DL, located at 571 Mbp – 637 Mbp. To test the hypothesis that this association was explained by the resistance gene *Bdv2* (located on chromosome 7DL), we investigated the haplotypes defined by the 16 SNP markers associated with BYD severity and were able to identify two haplotypes that exactly matched the presence or absence of *Bdv2* (Fig 4A). This same region was mapped using BYD severity and the presence or absence of *Bdv2* as a fixed covariate (Figure 4B). This analysis (Figure 4B) also detected a peak on chromosome 7AL. Lastly, we explored the effect of *Bdv2* on both BYD BLUEs and BLUPs, and we observed that the presence of *Bdv2* had a positive effect in reducing the disease severity by approximantly 10% (Figure 5A). The significant peak on chromosome 5AS, located at 46 Mbp – 103 Mbp, was explained by 10 SNP markers, comprising two main haplotypes, one of them associated with reduced BYD severity (Fig 5B). When we combined the different 5AS haplotypes with *Bdv2*, we observed that the presence of *Bdv2* had a positive effect, reducing the levels of BYD when combined with both 5AS haplotypes (Figure 5C), and suggesting an additive effect. Compared to the associations found for *Bdv2* (Fig 4B), we did not find any strong evidence of marker trait associations for the other evaluated traits (Figure S4).

**Figure 4.**
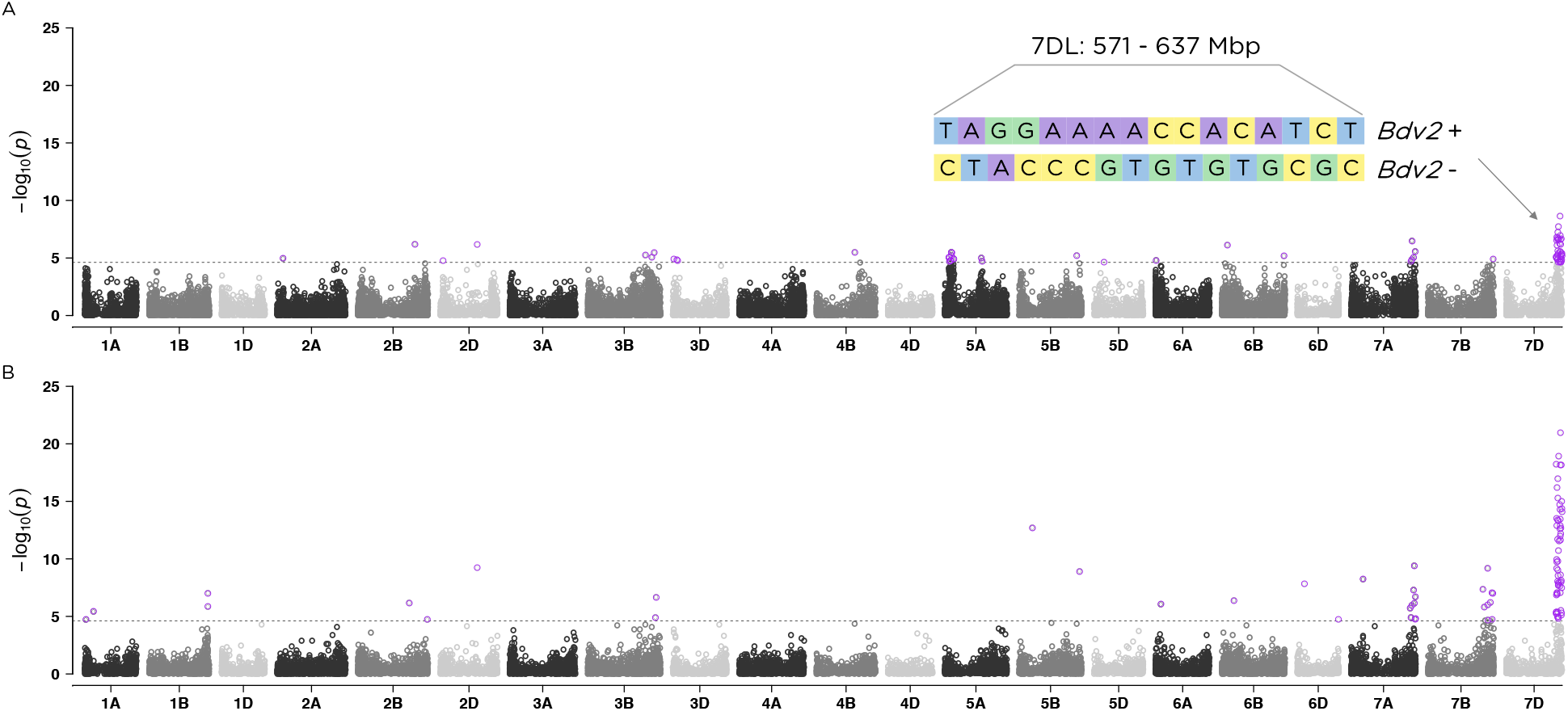
Manhattan plots showing the marker-trait associations using 346 wheat accessions and 29,480 SNP markers obtained with genotyping-by-sequencing (GBS) for A) BYD severity and B) presence/absence of *Bdv2* resistance gene. The 21 labeled wheat chromosomes with physical positions are on the x-axis and y-axis is the −log10 of the p-value for each SNP marker. Horizontal dashed lines represent the false discovery rate threshold at 0.01 level and data points highlighted in purple and above the threshold represent SNPs significantly associated with the trait. In panel a, the length of the region and the haplotypes defined by the significant SNP markers is displayed.

**Figure 5.**
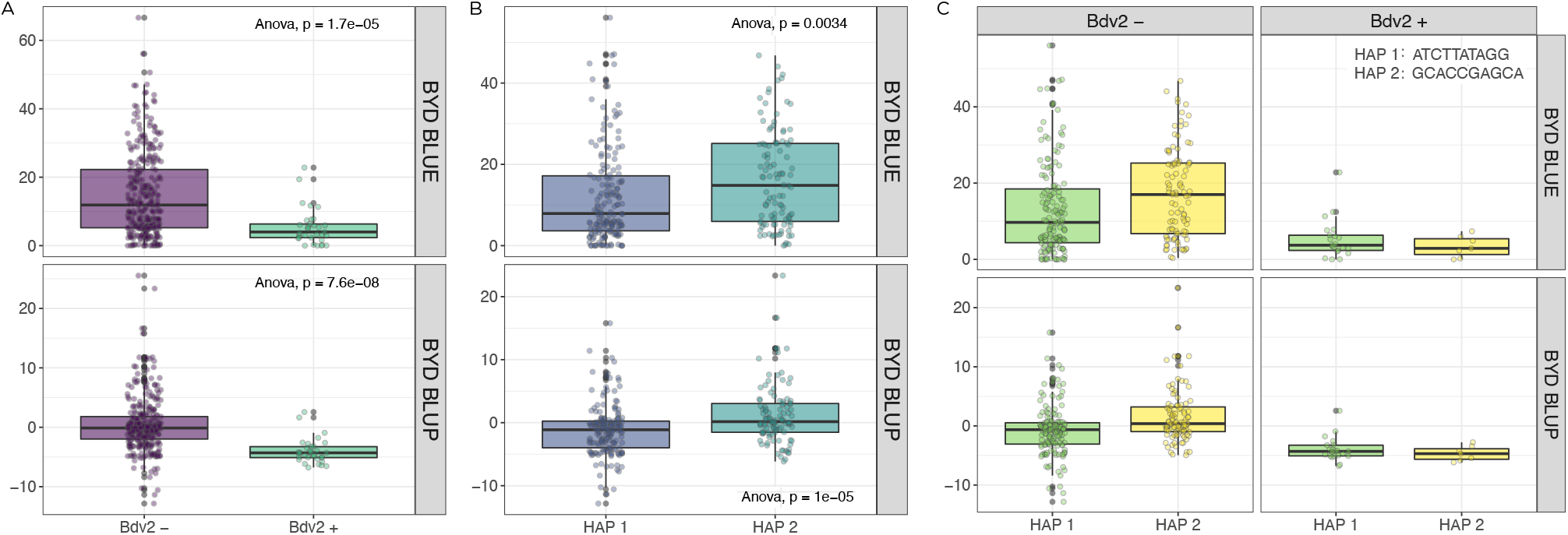
Measurement of barley yellow dwarf disease severity in wheat based on certain haplotype effects were panel A) represents the translocation segment carrying the resistance gene Bdv2, B) displays the two haplotypes for the significant region on chromosome 5AS, and C) shows the combination of Bdv2 resistance gene and 5A haplotype. Boxplots showing the significant reduction of BYD disease severity by averaging the phenotypic best linear unbiased estimated (BLUE) or best linear unbiased predicted (BLUP) values for the lines.

### Genomic Selection

To evaluate the potential of GS to predict BYD disease severity, we fit several GS models to the phenotypic BLUPs of BYD, PTH_M_, and reduction in GY. Across all traits, to determine predictive ability we used a five-fold cross validation where prediction ability ranged from −0.08-0.26. There was relatively good predictive ability for BYD severity ranging between 0.06 – 0.26, in comparison with PTHT_M_ and reduction in GY resulting in a lower range from 0.02 – 017 and −0.08 – 0.2, respectively (Figure 6). Evaluating the conformation of the training population, we observed that when including 2016-17 season, prediction abilities were the highest for BYD but the lowest for the other two traits, implying that season 2016 – 17 was either a good season to train the prediction models or a difficult season to predict based on available data.

**Figure 6.**
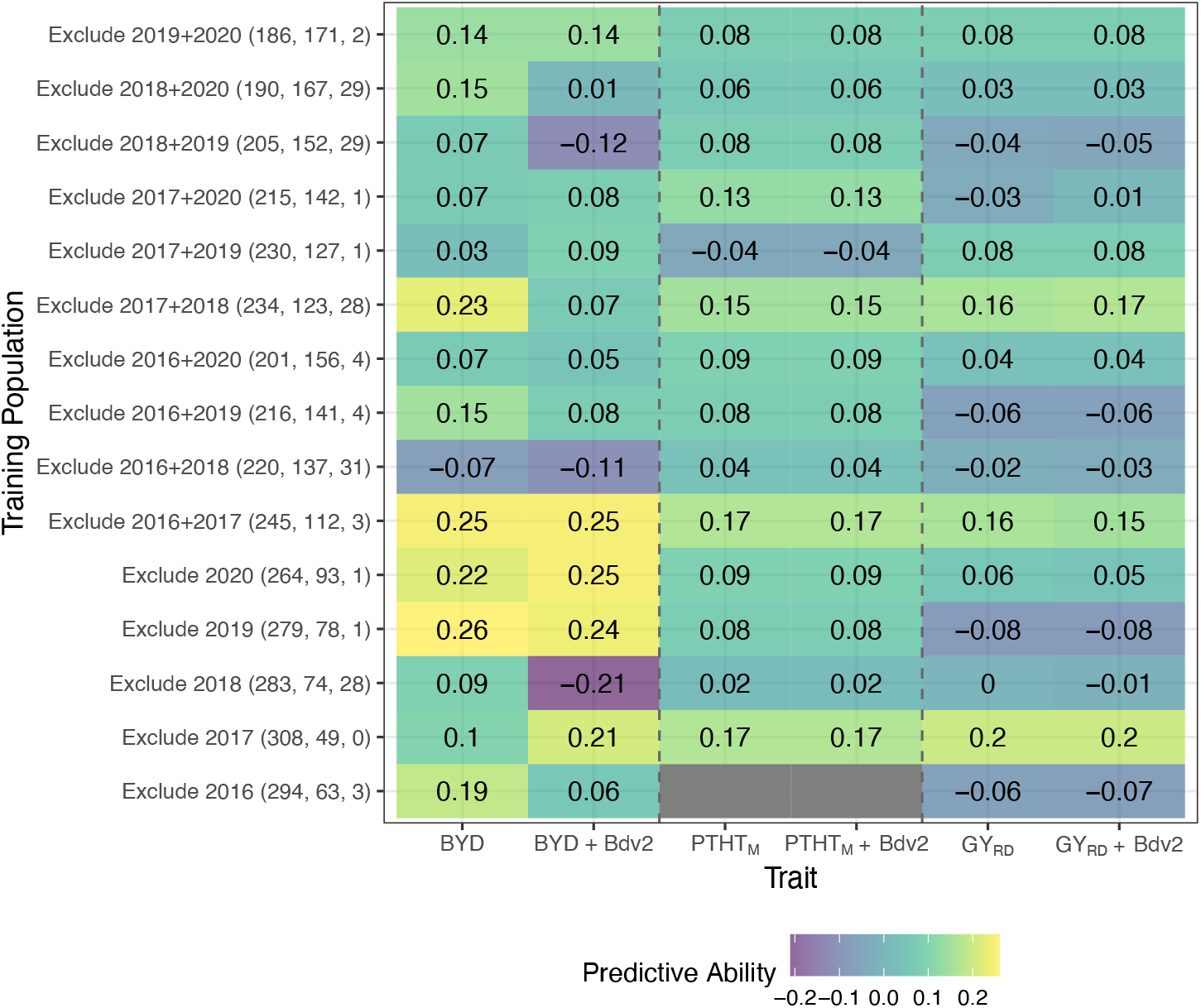
Genomic selection model predictive ability where each column represents one trait, and each row shows the conformation of the training population including size of training and testing population and number of lines with presence of *Bdv2* resistance gene. The value in each cell represents the predictive ability which is the correlation between the GS predicted value (GBLUP) and the phenotypic best linear unbiased predictor (BLUP).

To further investigate the power of GS, we developed models using a leave-two-out strategy, where two seasons were excluded from the training population and used as the testing population. We fitted GS models for all possible two-season combinations. This strategy resulted in slightly smaller training populations which decreased overall predictive ability (Figure 6). This result was evident for BYD predictions were excluding two seasons had a larger negative impact.

Lastly, we evaluated the effect of adding information about the genotype of the *Bdv2* resistance gene as a phenotypic fixed covariate into the GS models. There were differences in the effect of *Bdv2* on the predictive ability across BYD severity, PTHT_M_, and GY, showing a large effect for predicting BYD but almost no effect for PTHT_M_ and reduction in GY (Figure 6). The improved predictive ability for BYD was clearly reflected with the decrease of prediction ability obtained when season 2017 – 18 was excluded from the training population since most of the lines with the presence of *Bdv2* were evaluated in that season.

## Discussion

### Phenotypic Data

The success of breeding for BYD resistance is highly impacted by the ability to precisely characterize breeding material and disease symptoms. Even though BYD is spread worldwide, its incidence in a given year depends on several factors such as aphid pressure, planting date, and environmental conditions (e.g., temperature, rainfall, frost, etc.). In this study, we evaluated winter wheat advanced breeding lines during five seasons implementing a rigorous field-testing approach, that ultimately enabled us to consistently have plots contrasting with BYD infection and uninfected or low incident plots. Moreover, by using large yield-size plots we were able to calculate the reduction in GY and use this parameter as an estimate of field resistance.

The expression of BYD symptoms, however, was highly inconsistent during the different seasons. Seasons 2015-16 and 2016-17 showed the best expression of the disease symptoms, supported by the wide range of BYD severity between treated and untreated replications (Figure 1). Interestingly, both these seasons were conducted in the same experimental field (Table 1), suggesting that this location could favor the expression of BYD. Moreover, weather conditions were variable for all the seasons, suggesting that these had a huge impact on the disease occurrence. While temperature records were similar for all the seasons, precipitation records did show some differences. Season 2017 – 18 was dryer than normal, with 34% less precipitation than the 30 years historical average (1981 – 2010). On the other hand, season 2018-19 was wetter than normal, with 58% more precipitation than the 30 years historical average (Table S2).

### High-Throughput Phenotyping

Evaluating BYD resistance using visual phenotypic selection can be challenging due to the complex nature of the disease and rater variability (Poland and Nelson 2011). The use of HTP with UAS is gaining popularity within breeding programs because it further improves selection based on classical phenotyping. Accurate phenotyping is crucial for understanding the genetic basis of quantitative and complex traits like BYD. In this study, we used HTP to complement the visual BYD scoring. This tool improved our capacity for rapid, non-destructive, and non-biased evaluation of large field-scale numbers of entries for BYD resistance. We were able to determine strong correlation patterns between visual BYD severity and HTP derived parameters (Figure S3). However, none of the traits collected with UAS had a common genetic base with BYD severity (Figure 4 and Figure S4). Disease scoring using HTP is scaling fast among breeding programs; however, how to effectively use this data remains challenging. Some studies have shown that data collected with sensor-based tools can be substituted to improve classical disease visual evaluation (Sankaran *et al*. 2010; Kumar *et al*. 2016; Zheng *et al*. 2018); however, to the best of our knowledge this study is the first attempt to characterize BYD in wheat using HTP.

### Genome-Wide Association Analysis

Using GWAS we detected QTLs on chromosomes 5AS, 7AL, and 7DL for BYD severity BLUPs values. Using GBS tags that mapped to known alien fragments, we confirmed *Bdv2* resistance gene was located at 7DL, and confirmed that the 7DL QTL was explained by the presence of the *Bdv2* resistance gene. Even though only 33 wheat lines were positive for the presence of *Bdv2*, we still had enough power to detect its effect, suggesting that *Bdv2* has a strong effect on BYD under Kansas field conditions (Figure 5). The associations on chromosome 7AL, observed for both BYD severity and *Bdv2*, suggest that the SNP markers on the 7AL peak may be miss-anchored markers that should have mapped to 7DL. The relatively high heritability values obtained for the untreated replications (Figure 2) allowed us to detect a minor QTL on 5AS. Marza *et al*. (2005) reported a QTL at 38cM on the short arm of chromosome 5A associated with yellowing symptoms caused by BYD, and it is possible that this is the same region yet more data is needed to confirm if these QTLs are the same. The only other study reporting GWAS for BYD in wheat was able to identify several markers associated with BYD resistance on chromosomes 2A, 2B, 6A, and 7A (Choudhury *et al*. 2019b). However, most of the association were explained by individual SNP markers, and to date do not have any definitive biological link. GWAS results for the other traits used in this study did not discover genomic regions associated with the traits (Figure S4). Taken together, these results suggest that BYD resistance is not controlled by any large effect loci that could easily be incorporated into the breeding program, thus GS could be an efficient way to enhance BYD resistance.

### Genomic Selection

We evaluated several different GS models to identify the best approach for predicting BYD (Figure 6). Overall, we observed some trends including i) incorporating years with consistent BYD disease data in the training population increased the model predictive ability, ii) predicting years with high disease pressure is difficult, iii) using major effect QTL, such as *Bdv2*, had increased prediction performance, suggesting that it is responsible for much of the predictive power. These results suggest that GS based on G-BLUP with *Bdv2* as fixed effects would lead to the greatest genetic gain for BYD breeding. Using selected major QTL as a fixed effect to improve GS models was suggested in a simulation study (Bernardo 2014) and demonstrated with empirical studies (Rutkoski *et al*. 2014). Nonetheless, using *Bdv2* as a fixed effect in our GS strategies did not consistently improve the predictive ability for PTH_M_ or reduction in GY (Rice and Lipka 2019). However, there was not a consistent distribution of *Bdv2* allele across the cohorts. BYD predictions were low compared to other disease (reviewed by Poland and Rutkoski 2016). However, since this is the first report of GS for BYD resistance in wheat, we do not have similar results to make better comparisons. BYD has traditionally been reported to have low *H*^2^ (Tola and Kronstad 1984; Choudhury *et al*., 2019b) and in this study, even with well managemed plots that often had *H*^2^ approaching 0.8, we still had difficulty reproducing these results year to year as evidence of the challenge of studying this pathosystem. Moreover, the correlation between HTP parameters and BYD phenotypes was interesting, but not sufficient to be useful in combination with GS in the germplasm tested.

## Conclusions

We were able to show that *Bdv2* has a major effect controlling BYD resistance in the KSU breeding germplasm. Apart from the known *Bdv2* and a potentially novel 5AS region, we did not find evidence of other regions controlling BYD resistance supporting the hypothesis of limited resistance available in the current wheat gene pool and the highly polygenic nature of the trait. Moreover, our study was the first attempt to characterize and improve BYD field-phenotyping using HTP and apply GS to predict the disease. HTP traits showed strong correlation patterns with BYD severity, however, none of these parameters shared a common genetic architecture with BYD severity. The GS predictive ability results that we found in this study open the door for further improvement and testing GS implementation for breeding for BYD resistance. Continuing the improvement of BYD characterization and the search of new sources of resistance using species related to wheat, will be crucial to broadening the resistant genes available to introgress into wheat germplasm.

## Data Availability Statement

Supplemental material, including raw and analyzed phenotypic data, genotypic data, supplementary tables and figures, and basic plot scripts are available at <u>D</u>yrad doi:10.5061/dryad.ncjsxkswd (temporary link: https://datadryad.org/stash/share/xkKdr62QYB-YA93mkzq18_4yFUgwdnvv0uZUyuEHpAI) and GitHub https://github.com/umngao/wsm1_bdv2

## Supplementary data description

**Table S1 –** List of wheat entries phenotypically evaluated in the study. The table includes the type of entry (cultivar or breeding line), the season that the entry was evaluated, the result for the prediction of the presence/absence of the segment carrying the resistance gene *Bdv2*, and the best linear unbiased predictors (BLUPs) for all the phenotypic traits collected.

**Table S2 –** Precipitation (inches) during the five field seasons in Riley County, KS, where Rocky Ford and Ashland Bottoms experimental units are located. Normal temperature is defined as a 30–year average from 1981 – 2010. Data was obtained from Kansas State University (http://climate.k-state.edu/precip/county/)

**Figure S1 –** Growth trajectories and adjustment of the non-linear regression model of wheat lines for A-B) normalized difference vegetation index (NDVI) and C-D) digital plant height (meters). The data used correspond to season 2016 **–**17 phenotypic data. Calendar days is the number of days starting at January 1, 2017.

**Figure S2 –** Boxplots showing the phenotypic response of the wheat checks ‘Art’ (susceptible) and ‘Everest’ (tolerant) for A) barley yellow dwarf (BYD) disease severity (%), B) manual plant height (PTHT_M_) (m) and C) grain yield (GY) (tons/ha). Adjusted phenotypic values are shown for both insecticide treatment replications (treated and untreated).

**Figure S3 –** Scatterplots showing distribution and Pearson’s correlation values for the phenotypic traits studied during all the field seasons under two insecticide treatments (treated and untreated). A-B) season 2016 – 17, C-D) season 2017 – 18, E-F) season 2018 – 19, and G-H) season 2019 – 20.

**Figure S4 –** Manhattan plots showing genome-wide association analysis (GWAS) results for the phenotypic traits collected during the study.

## Acknowledgements

This material is based upon work supported by Kansas Wheat Commission Award No: B65336 “Integrative and Innovative Approaches to Diminish Barley Yellow Dwarf Epidemics Kansas Wheat”. PS was supported through a U.S. Fulbright-ANII Uruguay Scholarship. Any opinions, findings, and conclusions or recommendations expressed in this material are those of the author(s) and do not necessarily reflect the views of industry partners.

